# The thermostability of a Vaccine Delivery system X-Protein (VADEX-Pro) based protein nanoparticle

**DOI:** 10.1101/2023.02.15.528623

**Authors:** Gunn-Guang Liou, Ming-Chung Kan

## Abstract

We have adapted split GFP technology into the protein nanoparticle platform, Vaccine Delivery system X (VADEX), created in previous study. To evaluate the capability of this new platform, a model protein, maltose binding protein (MBP), was fused to the β-strand 11 of sfGFP and co-expressed with VADEX-10 which was composed of LYRRLE peptide and N-terminal part up to β-strand 10 of sfGFP. When these two fusion proteins were expressed in a cell, they were assembled into PNP spontaneously with a dynamic light scattering (DLS) particle size of 26 nm. This nanoparticle platform was renamed as VADEX-Pro for its capacity of expressing large protein on PNP. The thermostability of the assembled PNP was verified by both SDS-PAGE and DLS analysis following treatment. This PNP was stable at 25 °C and at temperatures as high as 40 °C for at least two months. Mutations that replaced cysteine residue of the LYRRLE peptide with serine or alanine destabilized and induced degradation of the VADEX-based PNP. The results in this study showed that the non-covalent complementation of split sfGFP became irreversible when reconstituted sfGFP was assembled in a VADEX-Pro PNP. This platform may be applied in developing thermostable vaccines.

## 1. Introduction

Fluorescent proteins have been identified in jellyfish, sea coral species and even arthropods [1]. These fluorescent proteins share one common beta-barrel structure in which 11 β-strands enclose an alpha-helix containing a three amino-acid chromophore. These β-strands provide structural support for precise spatial coordination required for chromophore maturation and function [2]. The fluorescent proteins have been applied in different fields; their applications include but are not limited to serving as a marker for cell biological study, an indicator for protein-protein interaction, and a monitor for proper folding of recombinant proteins [3]. Parts of the applications are based on split-GFP technology, in which these 11 β-strands of FP may be split into two or three uneven portions, and they can be either expressed individually or fused to two interacting proteins. And these split GFP portions can be reconstituted into a functional FP through strand complementation either through interacting between fusion partners or by following thermodynamic interactions [4,5]. When the GFP is split between strand 10 and 11, the complementation process is irreversible even under photo irradiation [6].

In our previous study, we created a novel self-adjuvanting protein nanoparticle (PNP) [7]. This PNP was assembled from a fusion protein containing two modules: a polymerization module and an antigen presentation module. The polymerization module is a modified amphipathic helical peptide derived from the M2 protein of type A influenza virus strain H5N1. And the antigen presentation module is a superfolder green fluorescent protein (sfGFP) that is able to incorporate peptide onto the surface of PNP through loop insertion. When this fusion protein was expressed in a cell, it self-assembled into PNP. We have identified a gain-of-function mutant that provides thermostability to the PNP based on the protein models built on transmission electronic microscope images. This technology is named VADEX-Pep, which stands for <u>Va</u>ccine <u>De</u>livery system <u>X</u>-<u>p</u>eptide. In this study, we have adapted split-GFP technology [5] into VADEX-Pep and created a new platform, VADEX-Pro (VADEX-<u>Pro</u>tein), and shown in our study results that we were able to incorporate large antigens onto the surface of VADEX-based PNP using sfGFP strand complementation. We evaluated the thermostability of the PNP assembled by VADEX-pro expressing MBP using both DLS and SDS-PAGE. These results suggested the incorporation of a large antigen through sfGFP strand complementation was irreversible when the sfGFP was assembled in a PNP. And the substitution of a single amino acid in the polymerization module induced PNP destabilization, whereas the PNP assembled by the original LYRRLE peptide was thermally stable in 40 °C incubation for two months.

## 2. Methods and materials

### 2.1. Plasmid construction

The VADEX-10 and strand 11 of sfGFP were synthesized by GeneArt. The nucleotide sequence of gene synthesized is listed in supplementary data. The VADEX-10 coding DNA was digested by NdeI and SalI and ligated with pET-Duet1 vector digested with NdeI and XhoI and the resulting plasmid was named VADEX-10. The strand 11 of sfGFP was digested by SalI and NotI and ligated with the VADEX-10 that had been digested by SalI and NotI; the resulting plasmid was named VADEX-Pro. The maltose binding protein coding region was PCR amplified from pD-MAL1 and inserted into the VADEX-Pro vector N-terminal in frame to the strand 11 or sfGFP between EcoRI and PstI. Site-directed mutagenesis of cysteine 8 was introduced by primers containing point mutations to the LYRRLE coding region.

### 2.2. Protein nanoparticle expression, purification and analysis

Protein expression and purification have been described in detail in a previous report. In short, the MBP-C PNP and its variants were transformed into the lipopolysaccharide defective ClearColi BL21(DE3) strain (Biosearch Technologies), and induced by 1 mM IPTG for 16-20 hours at 20 °C. After protein induction, bacterial lysate was generated by sonication and PNPs were purified using Ni-NTA resin and eluted by elution buffer (20 mM Na(PO_4_), 300 mM NaCl, and 500 mM Imidazole, pH 8.0), then stored at 4 °C. Before the thermostability test, the storage buffer of PNPs was changed from elution buffer to 0.5x storage buffer (10 mM Na(PO_4_), 150 mM NaCl pH 8.0), and the NaCl concentration was adjusted to a designated concentration using 5 M NaCl. DLS analysis was carried out in Stunner (Unchained Labs, Pleasanton, CA) using stunner plate (cat#7012025). For each sample being assayed, a triplicate of three micro-liters each were loaded onto microfluid channels in addition to a blank that using same buffer as sample. Protein concentration was determined by UV280nm and particle size was determined by the summary of four individual DLS readings. For the thermostability test, PNPs were incubated in a PCR block with continuous thermal control. Protein samples were taken out at designated time and denatured by mixing with 2x SDS sample buffer and heat denatured by 95 °C for 10 minutes before been stored in −20 °C freezer until ready for electrophoresis analysis. The negative stained TEM images of the MBP-C PNP were obtained by first cross-linking the purified MBP-C PNP (0.1 mg/ml) in freshly prepared fixation buffer (2% formaldehyde, 10 mM Na(PO_4_), 150 mM NaCl, pH 8.0) overnight. The protein samples were then processed for TEM by negative staining with uranyl acetate. Images were taken using a Tecnai G2 Spirit Twin.

## 3. Results

### 3.1. The design and expression of the VADEX-Pro PNP

The limitation on the size of peptide that can be inserted into the sfGFP loop poses a significant problem when developing subunit vaccines using the VADEX technology created in our previous study. In order to express large antigens on the surface of a VADEX-based nanoparticle, we have adapted split-GFP technology[5] into the VADEX platform. We fused the N-terminal of sfGFP, that contains β-strands 1-10 to the C-terminal of the polymerization peptide, LYRRLE, and generated the fusion protein, VADEX-10. The complementary β-strand 11 of sfGFP was fused to the C-terminal of a target antigen, the maltose binding protein (MBP) cloned from a protein expression vector, pD-MAL1, and the fusion protein is named MBP-11. These two fusion proteins were co-expressed in *E. coli* by constructing both ORFs in the same pET-Duet1 vector (Figure 1A). The PNP that was assembled by fusion proteins encoded in this vector was named MBP-C in this study. When we analyzed the protein expression profile that expressed only the target antigen, MBP-11, or both VADEX-10 and MBP-11, we found that co-expression of complementary fusion proteins was able to stabilize fusion proteins when comparing soluble fractions from bacteria expressing either single or both fusion proteins (Figure 1B). The his-tag for protein purification was fused only to the MBP-11 but not VADEX-10, so the only possibility that VADEX-10 can be purified by a Ni-NTA resin is because it was assembled into a PNP with the MBP-11 through sfGFP strand complementation (Figure 1B).

**Figure 1.**
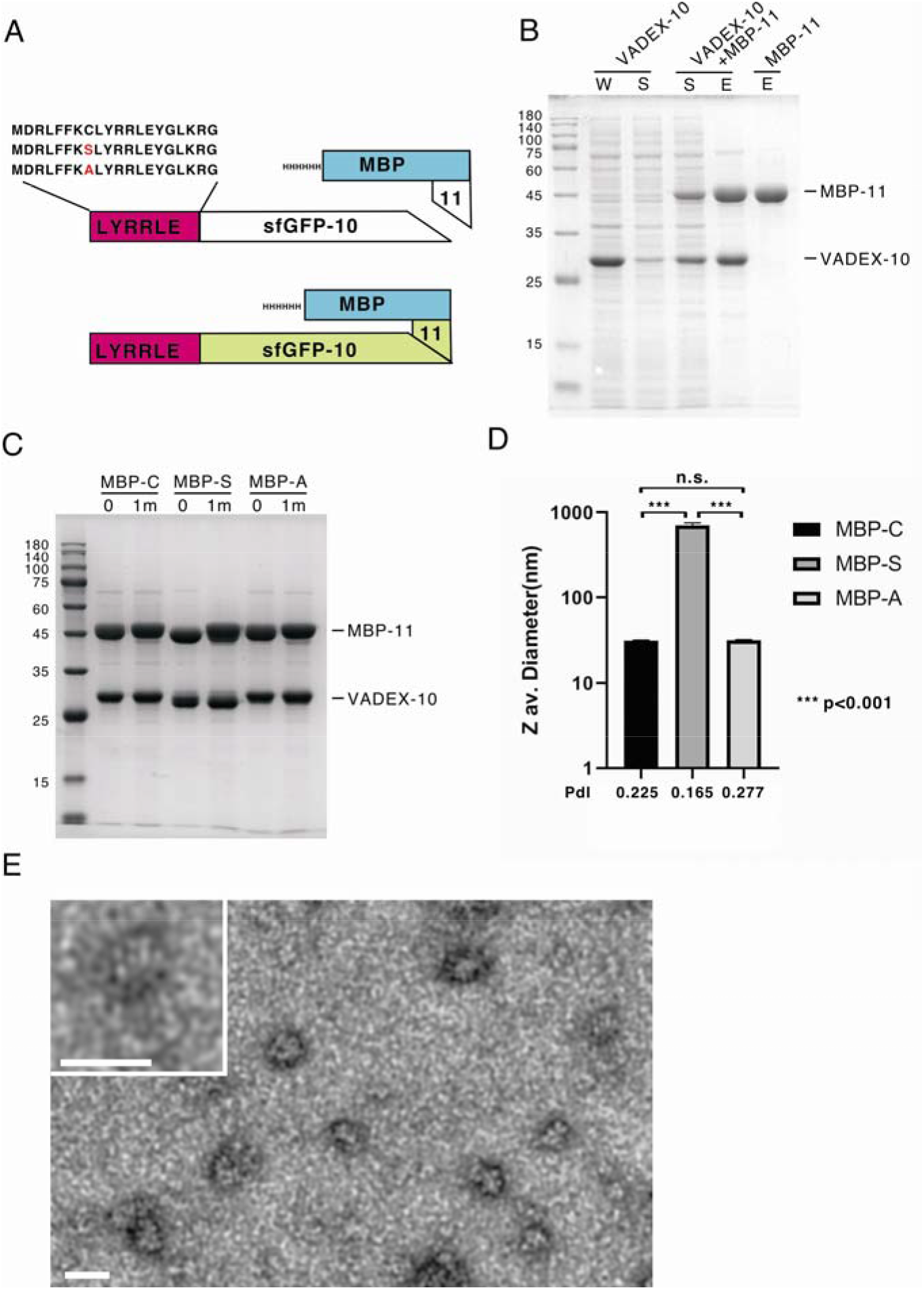
The design and protein expression of the VADEX-Pro protein nanoparticle and its variants. A) The N-terminal part of sfGFP containing β-strand 1-10 was fused to the C-terminal of the polymerization module, LYRRLE, and the fusion protein was named VADEX-10. The β-strand 11 of sfGFP was fused to the C-terminal of a model protein, maltose binding protein (MBP), and the fusion protein was named MBP-11. These two fusion proteins were co-expressed from the same expression vector, pET-Duet-1. The PNP that was assembled from these two fusion proteins was named MBP-C in this study. The cysteine residue in the MBP-C’s LYRRLE peptide was replaced by either serine or alanine and the PNP was renamed MBP-S or MBP-A, respectively. B) The whole cell lysate (W), soluble fraction (S), or eluted protein (E) from bacteria expressing either VADEX-10, MBP-11, or both fusion proteins were analyzed by SDS-PAGE. C) The purified MBP-C PNP and its variants, MBP-S and MBP-A PNPs, were analyzed by SDS-PAGE. The protein nanoparticles eluted from Ni-NTA resin were stored in elution buffer for one month before SDS-PAGE analysis to evaluate the stability of protein nanoparticles in elution buffer. D) The Z av. diameters (nm) in log scale of either MBP-C, MBP-S, or MBP-A PNPs in the storage buffer were analyzed by DLS in triplicate. Error bar represents the STDEV. The significance of the difference in particle size was determined by the student T test. (N=3) E) TEM image of MBP-C PNP after being fixed with 2% formaldehyde overnight. The size of the scale bar is 25 nm.

According to a protein model that was built on TEM images, the cysteine 8 (Cys8) residue in the polymerizing peptide, LYRRLE, has been predicted to mediate the formation of hydrogen bonds between two stacking dimers during the assembly of the VADEX protein nanoparticle. To test the role of the Cys8 in the assembly and stability of the VADEX-based protein nanoparticle, we made point mutations on the LYRRLE peptide by replacing the Cys8 with either serine (MBP-S) or alanine (MBP-A) in the MBP-C expression vector and examined the particle size and stability of these PNPs. The expression of MBP-C PNP reached 50 mg/L in flask culture, and the mutants also have similar expression levels (unpublished result). The VADEX PNPs that expressed MBP on the surface were stable in elution buffer at 5 °C storage for at least one month (Figure 1C). As determined by Dynamic Light Scattering (DLS), the particle size (Z av. diameter) of MBP-A was similar to that of MBP-C, whereas the particle size of MBP-S was much larger than either MBP-C or MBP-A. This excessive increase of MBP-S PNP particle size implied the substitution disrupted the assembly process and resulted in a non-specific protein aggregation. Although the average particle size was similar between MBP-C and MBP-A, the polydispersity index (PdI) of MBP-A was significantly higher than MBP-C, which suggested particle destabilization and a drastic increase in particle size variation due to Cys-to-Ala substitution (Figure 1D).

To verify whether MBP-C still shares the same geometry of VADEX-Pep PNP[7], we crosslinked the purified MBP-C PNP and then examined the PNP structure under TEM after negative staining. The MBP-C PNP still maintained similar geometry to VADEX-Pep as shown in previous studies, only with a shorter long axis which is consistent with the results from DLS analysis. From DLS analysis, the Z av. diameter of fresh prepared VADEX-Pep (LYRRLE-sfGFP-2xhM2e) was between 38-40 nm whereas that of MBP-C was 28 nm. This difference in particle length could be attributed to the large molecular size of MBP. With a molecular weight of 45 kDa, it is about twice the MW of the sfGFP or six folds the MW of the inserted loop on VADEX-Pep. The bulky MBP attached to the distal end of sfGFP most likely exceeded the space limit for each subunit, pushing sfGFP and its cargo toward both ends and forming the wine barrel shaped particle as seen in the magnified image (Figure 1E).

### 3.2. The stability of VADEX-Pro PNP at 25 °C storage

We want to evaluate the stability of VADEX-Pro PNP variants when they are stored in physiological buffer at 25 °C. The stability of these proteins was determined by both SDS-PAGE and DLS for the integrity of PNPs that remained soluble. When the storage buffer was replaced with phosphate buffer (pH 8.0) with 150 mM NaCl, the MBP-S PNP became unstable at 25 °C and the MBP-11 was either degraded or precipitated within the first two weeks. The MBP-A PNP was more stable than the MBP-S PNP, but it was hydrolyzed into a smaller size during the second month. The MBP-C PNP was stable for more than one month before showing signs of protein hydrolysis (Figure 2A). The stability of MBP-C PNP was also supported by the results of the DLS analysis. The Z average diameter of PNP assembled by wildtype peptide (MBP-C) was 26.36 nm and was shown as a single peak (Figure 2B, left panel). Whereas the DLS profile of the PNP assembled by peptide contains a Cys-to-Ala substitution (MBP-A) and was split into two peaks of different sizes (Figure 2C, left panel). The first peak of MBP-A matched with the DLS profile of MBP-11 (Figure 2D, right panel) suggests the purified MBP-A PNP contains two populations, the assembled PNP and dissociated free MBP-11 proteins. After being incubated at 25 °C for 2 months, the MBP-C particle remains intact with increased variation in particle size (Figure 2B, right panel). For MBP-A PNP, the percentage of dissociated MBP-11 protein has increased from 55.2% to 71.1% after 2 months of storage at 25 °C (Figure 2C, right panel). The results from DLS analysis of MBP-C and MBP-A PNPs are consistent with the results from SDS-PAGE and we concluded that the MBP-C PNP has high stability and can sustain long-term storage at 25 °C, and the substitution of Cys8 by alanine decreases PNP stability.

**Figure 2.**
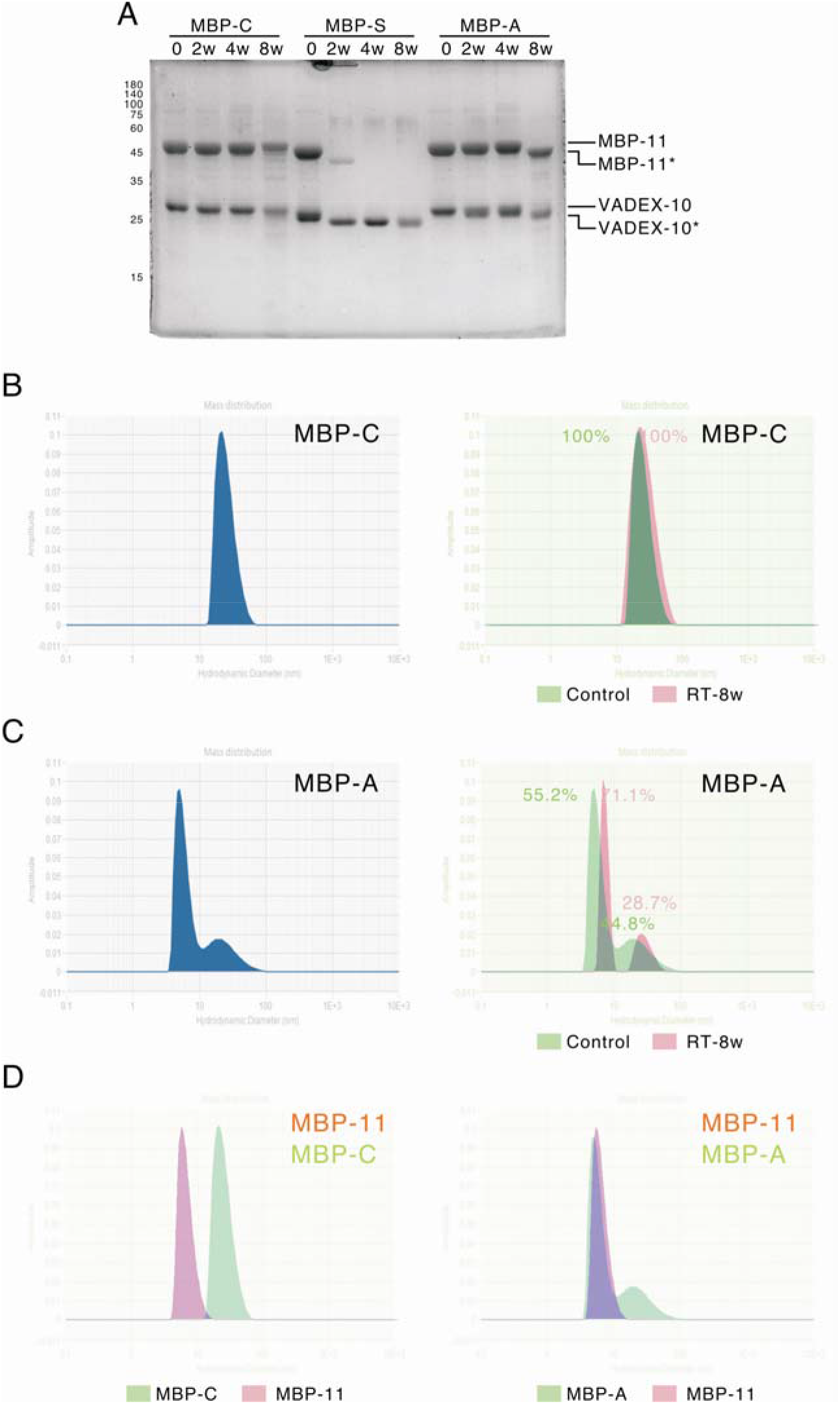
The stability of MBP-C and its variants at 25 °C storage in a physiological buffer. A) MBP-C PNP and its variants were stored at 25 °C for up to two months before protein samples were taken for SDS-PAGE analysis of protein integrity. The hydrolyzed MBP-11 and VADEX-10 fusion proteins were marked by an asterisk “*”. B) The stability of MBP-C PNPs was analyzed by DLS for particle stability at 25 °C storage. The left panel was fresh prepared control MBP-C PNP in a phosphate buffer containing 150 mM NaCl. The right panels were the result after merging left panel [10] with DLS result from MBP-C PNP that had been stored for 2 months at 25 °C (pink). C) The same DLS analysis results of MBP-A PNP were shown. D) The comparisons of MBP-C (left panel) and MBP-A (right panel) PNP DLS polygraphs to that of MBP-11 were shown.

### 3.3. The stability of VADEX-Pro PNP in high temperatures storage

The results from figure 2 highlighted the importance of Cys8 in LYRRLE peptide mediated PNP assembly. To test the thermostability of MBP-C and MBP-A PNPs at higher temperatures, we incubated both PNPs at a series of elevated temperatures for one hour and then verified the structural integrity of sfGFP by fluorescence imaging and particle stability by DLS analysis. Both MBP-C and MBP-A PNP lost fluorescence when the temperature reached 80 °C (Figure 3A), which is consistent with previous reports on the thermostability of split-GFP and sfGFP [8,9]. The Z average diameter of MBP-C PNP increased by 50% when the temperature was elevated from 40 °C to 50 °C, whereas the particle size of MBP-A PNP increased by three folds under the same conditions (Figure 3B). This data suggested that both sample’s particle sizes peaked at 70 °C. Considering the apparent stability of MBP-C at 25 °C and particle integrity after storage, we decided to focus on MBP-C when testing the long-term thermostability of VADEX-based PNP. In a previous study, the salt concentration in the storage buffer was found to affect the stability of VADEX-based PNP [7]. To test the effects of NaCl concentration on MBP-C PNP long-term thermostability, the NaCl concentration in the storage buffer was adjusted to either 300 mM, 600 mM, or 900 mM, and PNPs were then maintained at either 40 °C or 50 °C and followed for 8 weeks. The thermostability of PNP was monitored in three categories: the protein integrity was evaluated by SDS-PAGE; the particle size was determined by DLS; and the concentration of PNP that remained in its soluble form was measured by UV 280 nm.

**Figure 3.**
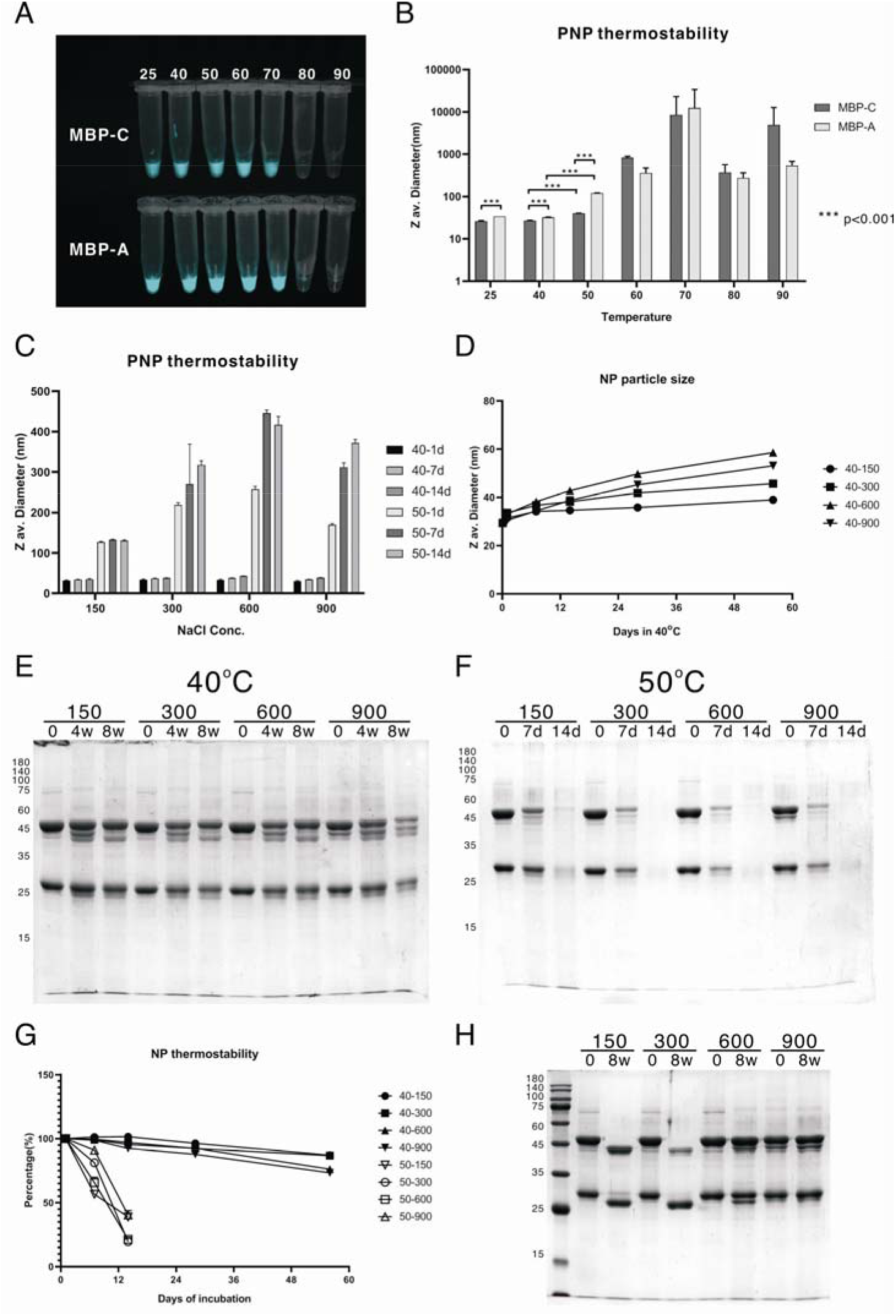
The thermostability of MBP-C and MBP-A protein nanoparticles. A) The fluorescence of MBP-C and MBP-A PNPs after being incubated at a series of elevated temperatures for one hour. B) The Z average diameter (Z av. diameter) of MBP-C and MBP-A PNPs after being incubated in different temperatures for one hour were compared. (N=3) C) The Z av. diameter of MBP-C PNPs incubated at 40 °C or 50 °C for a period up to 14 days in storage buffers containing different NaCl concentrations was shown. (N=3) D) The hydrodynamic particles size of protein nanoparticle incubated in phosphate buffer containing either 150 mM, 300 mM, 600 mM, or 900 mM NaCl at 40 °C was followed for 8 weeks. (N=3) E) The integrity of protein nanoparticles stored in phosphate buffer containing 150 mM, 300 mM, 600 mM, or 900 mM NaCl at 40 °C was followed for a period up to 8 weeks. The protein samples were collected at designated time points and stored at −20 °C before being analyzed by SDS-PAGE. F) The same experimental design as in Fig. 3E, except the storage temperature was 50 °C. G) The concentration of protein nanoparticle remained in solution after being stored at either 40 °C or 50 °C was monitored for 8 weeks. (N=3) H) The MBP-C PNP in phosphate buffer containing different concentrations of NaCl was stored at 4 °C for 2 months, and the protein integrity was analyzed by SDS-PAGE. The presented data is the representative result of two independent experiments.

When the MBP-C PNP was incubated at 40 °C, the particle size was relatively stable with an increase in the Z average diameter ranging between 24.4 nm (150 mM NaCl) to 49.7 nm (600 mM NaCl) after 8 weeks. But when the PNP was incubated at 50 °C, the Z av. diameter of MBP-C PNP increased dramatically within one day by 3.3 to 7.7 folds (Figure 3C). This dramatic increase in particle size was correlated with the precipitation and removal of PNP from solution within two weeks (Figures 3F, 3G). These data suggested the MBP-C PNPs were relatively stable at 40 °C storage for at least two months without showing significant PNP particle size change or protein lost to denaturation in a phosphate buffer without protein stabilizing excipients (Figure 3E). And the PNP particle size and protein integrity were not affected by salt concentration when PNP was incubated at 40 °C (Figure 3E). Whereas when these PNPs were stored at 50 °C, the half-life was less than 14 days due to protein denaturation and precipitation, a result that was similar for MBP-C PNPs in different NaCl concentrations (Figure 3G). In contrast, after two months storage at 4 °C in buffers containing either 150 mM or 300 mM NaCl, both VADEX-10 and MBP-11 were hydrolyzed into sMAL1er fragments (marked with asterisk). The MBP-C PNP stored in buffers containing 600 mM or 900 mM NaCl was more stable and only partially hydrolyzed (Figure 3H). This result suggests that the MBP-C PNP degradation at 4 °C and 50 °C is through different pathways. However, the results are consistent with previous studies that VADEX-based protein nanoparticle was stabilized by the presence of high salt concentration that strengthened hydrophobic interactions within the PNP [7].

## 4. Discussion

In this study, we have shown that a large protein like MBP, can be incorporated onto the VADEX-based PNP through sfGFP strand complementation. The assembled PNP is stable under long-term storage at ambient temperatures up to 40 °C and the reconstituted sfGFP can withstand high-temperature incubation at the same level as full length sfGFP. The complementation between strand 11 and the rest of GFP was reported as irreversible even under the condition of photo-irradiation[6]. Our data also support this conclusion, since the DLS polygraph from MBP-C PNP after 2 months storage at 25 °C showed no sign of dissociation. In contrast, when the integrity of MBP-C PNP was weakened by the substitution of Cys8 by alanine, the reconstituted sfGFP fell apart and the amount of dissociated antigen increased over time, as expected for proteins bound by non-covalent strand complementation.

The MBP-C PNPs aggregated non-specifically into large particles and precipitated in a salt-independent manner when stored at 50 °C. This salt-independent aggregation may be due to the partial denaturation of MBP, which has a Tm of 62.3 °C at pH 8.2 in the absence of maltose [11]. The overall stability of a PNP depends on the least stable component of the particle. In our previous studies, the unstructured peptide antigen (hM2e) became the target of water hydrolysis even when the particle size remained stable [7]. Although other factors besides structural integrity, like zeta-potential or salt strength, are also important for the stability of PNP at high temperatures [12]. The MBP used in this study was cloned from the pD-MAL1 vector with an isoelectric point (pI) of pH 4.9, which will create a large zeta-potential for preventing PNP aggregation in the phosphate buffer (pH 8.0) used in this study. The hydrolysis of MBP-C PNP in 4 °C storage under salt concentration dependency is consistent with previous studies [7], This data suggested that the storage at low temperature may alter the structure of LYRRLE peptide and reduce its ability to assemble tightly into PNP, a caveat that can be compensated for by using buffers with a higher NaCl concentration.

In the previous study, Cys8 in the LYRRLE peptide was predicted to form a hydrogen bond with the glutamic acid from another dimer and stabilize the PNP [7]. In this study, we provided additional evidences proving the importance of Cys8 in VADEX PNP stability. The cysteine in a protein structure plays dual roles depending on its location; it forms part of the hydrophobic core when it is buried inside a protein, a role that may be substituted by alanine. Or, cysteine may form either a hydrogen bond or a disulfide bond within or between proteins through its strong nucleophilic property when it is exposed to the surface, a role that can be partially replaced by serine. In this study, the PNP maintained similar size but less stability after replacing the Cys8 with alanine, but the PNP particle structure and stability were greatly compromised by replacing the Cys8 by serine. These results suggest the Cys8 in the LYRRLE peptide serves mainly as part of the hydrophobic core during the PNP assembly, yet the nucleophilic property of Cys8 also provides higher stability to the PNP, likely by forming a hydrogen bond with Glu14 as predicted by protein modeling.

The thermostability of several PNP platforms has been evaluated in various studies. The PNP platform based on hyperthermophilic bacterial aldolase has been shown to be able to withstand short-term (one hour) temperature incubation up to 75 °C, in which 80 % of the proteins remained soluble post treatment [13]. In another study, an artificially evolved PNP, I53-50, when assembled with the receptor binding domains (RBDs) of SARS-CoV2 spike protein that had been stabilized by Deep Mutational Scanning, was able to maintain particle integrity after being incubated for 28 days at 37-40 °C [14]. But the PNPs being evaluated in these thermostability studies were not immunogenic; they have to be mixed with oil-in-water type adjuvants before immunization for activity [13,15–17] and the thermostability of these PNPs in their final formulation has not been reported. To gain thermostability after being formulated with an oil-in-water adjuvant, these PNPs may need additional processing like lyophilization or spray drying [18,19]. In comparison, VADEX based PNP is able to activate humoral immune responses without additional adjuvant [7] and it is a suitable platform for developing readily injectable therMAL1y stable vaccines.

## 5. Conclusions

In conclusion, we have adapted split-GFP technology in VADEX PNP and enabled the incorporation of large proteins onto the surface of VADEX PNP through irreversible strand complementation, and this self-adjuvanting platform may serve as a basis for developing therMAL1y stable vaccines.

## Institutional Review Board Statement

This study did not perform experiments that required approval from institutional review board.

## Informed Consent Statement

No informed consent statement required in this study.

## Data Availability Statement

The data will be available upon request.

## Acknowledgments

We thank Image Core of Institute of Molecular Biology at Academia Sinica for kindly providing an FEI Tecnai G2 spirit EM to be used in this study.

